# Winter-run Chinook salmon juvenile recruitment and early life history response to flow in a heavily altered tailwater

**DOI:** 10.64898/2026.09.24.754146

**Authors:** Chase A. Ehlo, Brad Cavallo, Kai Ross, Steve Zeug

## Abstract

Diverse life-history strategies allow species to spread risk across a mosaic of habitat conditions. In Chinook Salmon (*Oncorhynchus tshawytscha*), particularly in the Central Valley of California, this is expressed through varied adult run timings and juvenile outmigration strategies. Anthropogenic impacts, particularly dams, disrupt these fundamental life-stage transitions, often forcing managed populations to adapt to homogenized, less stochastic hydrologic regimes. This is exemplified in the Sacramento River, where the conservation of endangered Sacramento River winter-run Chinook salmon requires balancing complex water operations with the ecological needs of a population confined to a short stretch of suitable habitat downstream of Shasta Reservoir. Using a 23-year dataset (2002–2024), we evaluated the combined effects of flow, temperature, and spawner abundance on juvenile production and life-history expression in this habitat. We found that flow and spawner abundance best predicted juvenile abundance at Red Bluff Diversion Dam while temperature had considerably less support. The proportion of juveniles migrating as smolts exhibited significant density dependence that was negatively correlated to both female spawner abundance and peak flows. These results suggest focusing on temperature management alone may be insufficient. Effective management strategies should consider flow variability and habitat restoration to facilitate varying migration strategies and expand upstream rearing capacity. By addressing these physical and hydrologic constraints, managers can better support the full suite of life-history strategies necessary for the resilience of winter-run Chinook salmon.

## INTRODUCTION

Pacific salmon (*Oncorhynchus* spp.) are characterized by an expansive life cycle that spans freshwater natal streams, estuarine corridors, and marine feeding grounds over the course of 2 to 8 years before culminating in a return to natal tributaries for reproduction (Groot and Margolis 1991; Crozier et al. 2008). Consequently, population declines present a major challenge for conservation and management, as this variable spatial and temporal scale can complicate identification of the life stages being impacted and the factors limiting those life stages (Kendall et al. 2023). In lotic systems where spawning, egg incubation, and juvenile rearing occur, large-scale alterations resulting from dam construction, flow regulation, hatchery operations, and land use changes can individually and synergistically narrow opportunities for successful life stage transitions and result in impacts to populations (Merz et al. 2024).

This is most evident in the highly contentious Central Valley in California where rivers support four unique run timings of Chinook Salmon (*Oncorhynchus tshawytscha*) named for the season of entry into freshwater (winter, spring, fall, and late-fall; (Yoshiyama et al. 1998). In addition to variation in adult migration timing, the different runs also vary in the spatial distribution of spawning habitat, and time spent in freshwater before reproduction. Juvenile Chinook Salmon express multiple migrations strategies that spread risk and take advantage of the mosaic of habitats that activate under different hydrological conditions. Juveniles may disperse from natal habitat soon after hatch and subsequently rear in floodplains, tributaries, or the estuary prior to ocean entry. Or they may remain in natal habitat and rear until smolting, initiating migration as young-of-the-year (ocean type) or yearlings (stream type). This plasticity promotes life history diversity and bolsters population resilience (Carlson and Satterthwaite 2011; Satterthwaite et al. 2017; Phillis et al. 2018).

Sacramento River winter-run Chinook Salmon are currently listed as endangered under the Federal Endangered Species Act (Register 1994). Historical spawning habitat for this run included much of the upstream reaches of the Sacramento River, including major tributaries such as the McCloud and Pit Rivers, where holding and spawning conditions, particularly water temperature, were favorable during the summer spawning season (Williams 2006). Access to these historical spawning grounds was blocked by the construction of Shasta (1945) and Keswick (1950) Dams, and a single population is now confined to a 40 km reach of river in the valley floor downstream of Keswick Dam far outside of their historical spawning range (Williams 2006). This restricted reach of the Sacramento River limits both the recovery potential of the run and the operational flexibility of the Central Valley’s water infrastructure. This is most evident during critical incubation and emergence periods, where Shasta and Keswick Dams are managed to minimize early life-stage mortality, a task increasingly complicated by the hydrologic uncertainties of a changing climate (Nickel et al. 2004; Yates et al. 2008; Beer and Anderson 2013; Sapin et al. 2017; Hallnan et al. 2020).

Current management is almost entirely focused on water temperature and its effect on embryo survival (U.S. Fish and Wildlife Service 1999; Martin et al. 2020; Anderson et al. 2022). While temperature can be a strong driver of mortality, particularly in drought years, the relative influence of other stressors, including flow volume, physical habitat quantity and quality, and predation, remains poorly quantified (NASEM 2026). Density-dependent effects may further complicate early life stage dynamics in this confined spawning and rearing habitat for winter-run Chinook Salmon. These may range from the spawning stage where females are competing for space to construct redds, to resource competition among rearing juveniles which can be further compounded by environmental stressors (Peterson et al. 2020). In turn, this may decrease upstream rearing and increase migration of fry and parr into lowland river reaches (Greene and Beechie 2004; Williams 2006). Habitats in the lowland reaches have often been viewed as hostile for juvenile salmon, due to proliferation of non-native predators and channelization that limits habitat utility (Michel et al. 2015; Hellmair et al. 2018). However, when high discharge periods overlap with suitable environmental conditions, juveniles migrating downstream can access productive floodplains, tidal wetlands, and non-natal tributaries that provide high-growth rearing alternatives that increase rearing capacity (Jeffres et al. 2008; Katz et al. 2017; Phillis et al. 2018).

Collectively, the interaction between environmental drivers, juvenile production, and density-dependent population dynamics remains poorly resolved within this spatially restricted reach, thereby limiting conservation, restoration, and operational decisions. To begin addressing this deficiency, we leveraged a 22-year monitoring dataset to provide inference about the roles of environmental drivers, and density dependence in shaping juvenile production and migratory life-history expression of this endangered run. We pursue this through three separate but complementary analyses:

1. Modeling total juvenile production as a function of environmental and biological covariates
2. Modeling the environmental and biological drivers of juvenile life history expression.
3. Estimating the potential for density dependent production of different juvenile migratory phenotypes.

The results of this analysis provide new insights into population dynamics of this endangered run that can be used to address stressors that are likely to be limiting for certain life stages and adjust management decisions to support juvenile productivity within the constraints of current habitat.

## Methods

### Study Area

The Sacramento-San Joaquin River system represents one of the most heavily altered hydrologic landscapes in the world, requiring a delicate balance between an extensive agricultural economy, the water needs of densely populated metropolitan centers, and the conservation of native flora and fauna. The Sacramento River is the largest in California and drains portions of the Southern Cascades, Sierra Nevada, and Coastal Range. As discussed earlier, the river is impounded by Shasta Dam that was completed in 1945. Downstream of Shasta Dam is a smaller reservoir impounded by Keswick Dam that smooths out discharge fluctuations from Shasta Dam operations and serves as the upstream extent of anadromous fish migration. From Keswick Dam the river flows south for approximately 486 km before entering the upper reaches of San Francisco Bay. Our study focused on the approximate 100 km section of the Sacramento River from Keswick Dam to the decommissioned Red Bluff Diversion Dam where a juvenile monitoring station is operated. Most of the winter-run spawning occurs in the first 9 km of the study reach, but, in the past, extend downstream almost 50 km to the confluence with Battle Creek (Fig. 11). California’s Mediterranean climate brings cool, wet winters and warm, dry summers. However, Shasta Dam operations significantly alter the Sacramento River’s natural seasonal flow pattern (Zeug et al. 2011). Peak flows are now attenuated during the wet season to store water for release during summer, when agricultural and urban demands peak (Nichols et al. 1986). To better manage the reservoir’s cold water volume needed for winter-run spawning temperature requirements while also providing for power generation, a temperature control device (TCD) was installed on Shasta Dam in 1997. This device can access and blend water from different depths while maintaining the hydropower capacity of Shasta Dam (Hanna et al. 1999).

### Data collection

We collated 23 years of adult and juvenile winter-run monitoring data for this analysis (2002-2024). We began with 2002 because no juvenile data were collected during 2000-2001, and methodologies for adult surveys were not standardized prior to 2000. The California Department of Fish and Wildlife (CDFW) and U.S. Fish and Wildlife Service coordinate to complete adult population estimates via carcass surveys. They estimate spawner abundance by marking carcasses during weekly surveys and applying a Cormac Jolly-Seber mark-recapture model to estimate abundance. Detailed methodology of these surveys are reported in Killam (2025).

Migrating juvenile Chinook Salmon are monitored by the US Fish and Wildlife Service at the decommissioned Red Bluff Diversion Dam (RBDD) located approximately 94 km downstream from Keswick Dam (Fig. 1). Detailed methodology for this sampling program and methods can be found in Poytress et al. (2014). In short, passage is estimated by extrapolating captures of juvenile Chinook Salmon in multiple rotary screw traps (RSTs) located immediately downstream of the RBDD. Using capture efficiency data from mark-recapture trials, raw catch data are expanded to an estimate of total passage. Further, Chinook Salmon migrate at different life stages and these are estimated separately based on size, where fish ≤ 45mm are classified as *fry* and fish > 45 mm are classified as *pre smolts/smolts*. The data for both life stages are combined into a single value of *fry equivalents* by applying a fry-to-smolt survival value. This value is colloquially referred to as the Juvenile Production Index or JPI which is used in regulatory frameworks (Poytress and Mccraney 2025). For this analysis we use passage estimates for the two different migratory life histories as well as the combined JPI. Because rotary screw traps classify *fry* and *pre-smolts/smolts* solely by size, their actual life history stages likely overlap. To account for this, we refer to fry-sized fish as *non-natal rearing* and smolt-sized as *natal rearing*. *Non-natal* rearers are described by Apgar et al. (2021) as an early life-stage variant that hatches and quickly departs the natal stream, whereas *natal* rearers utilize their birthplace habitat for up to six months.

**Fig. 1.**
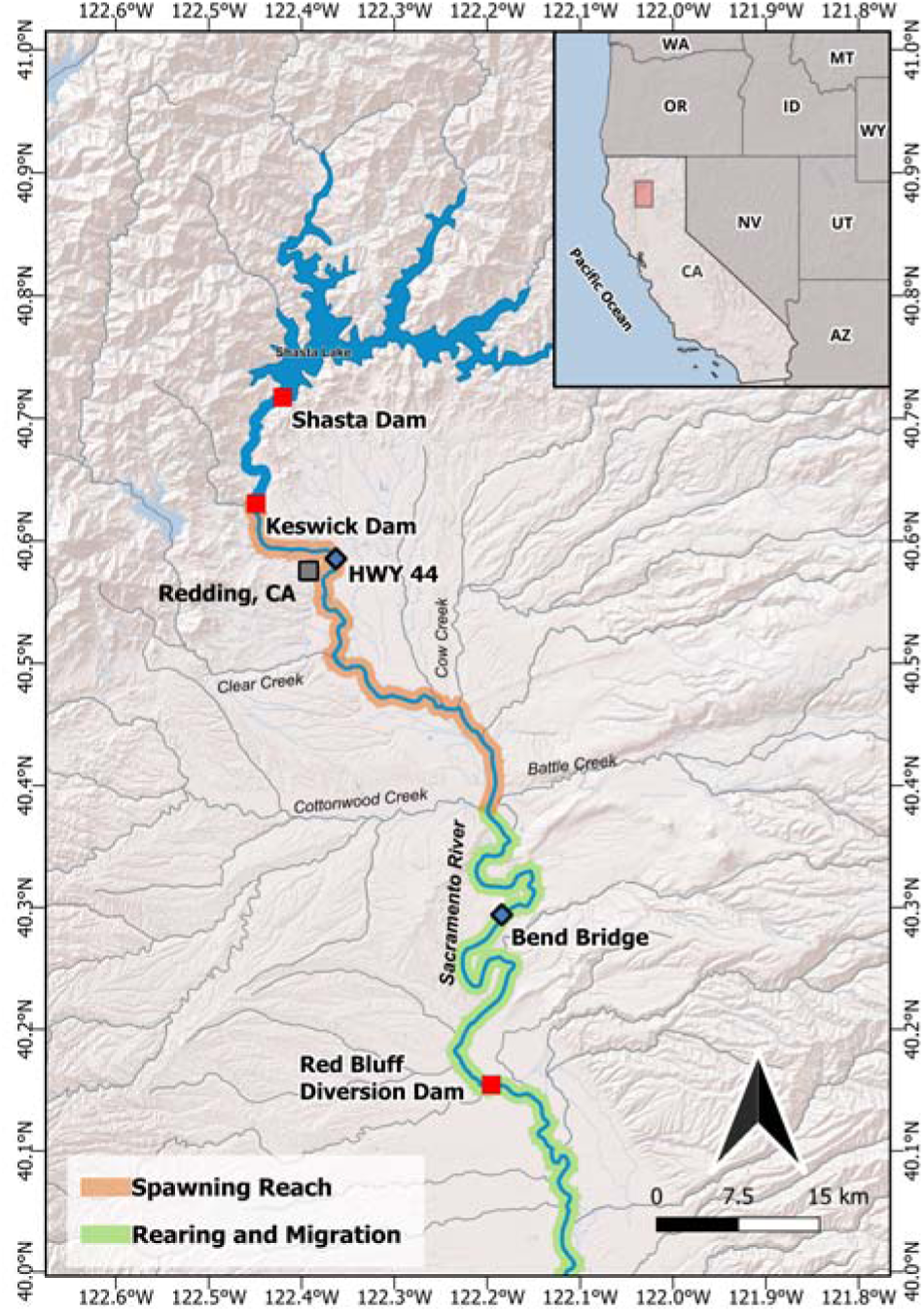
Map of the study area showing approximate spawning (orange) and migratory (green) reaches of winter-run Chinook Salmon in the Sacramento River from Keswick Dam downstream to Red Bluff Diversion Dam, CA.

### Environmental Data Collection

We selected environmental variables based on hypothesized drivers of variation in juvenile production and life history expression, including both flow and temperature data (Table 1 and Table S1 in supplementary materials). We considered three temperature variables with the first two metrics calculated at a station near the California State Highway 44 bridge (Fig. 1). We selected this location because it lies near the midpoint of winter-run spawning habitat. For time periods when data were unavailable at this station, we interpolated values using a linear model with data from an upstream station just below Keswick Dam and a station 9 km downstream near the limit of current winter-run spawning (Fig. 1).

**Table 1.** Summary statistics for environmental covariates used in JPI and natal rearing proportional passage modeling.

| Variable | Units | Months | Mean (± SE) |
| --- | --- | --- | --- |
| Females<sup>1</sup> | Individuals | May-September | 2985 ($\pm 525.25$) |
| Temp_SAC_I<sup>1</sup> | °C | July-September | 11.75 ($\pm 0.17$) |
| DD_above_12.0_I<sup>1</sup> | °C | July-September | 23.64 ($\pm 8.81$) |
| Temp_RB_M<sup>1</sup> | °C | October | 13.62 ($\pm 0.23$) |
| Flow_IE<sup>1</sup> | m<sup>3</sup>/sec | May-October | 308.87 ($\pm 11.98$) |
| Total_discharge_IE<sup>1</sup> | m<sup>3</sup>/sec | May-October | 4.9 x 10<sup>9</sup> ($\pm 2.0$ x 10<sup>8</sup>) |
| Flow_M<sup>1</sup> | m<sup>3</sup>/sec | October | 217.22 ($\pm 5.95$) |
| Flow_CV_M<sup>1</sup> | - | October | 0.08 ($\pm 0.01$) |
| max_flow<sup>2</sup> | m<sup>3</sup>/sec | October | 213.45 ($\pm 6.47$) |

<sup>1</sup>Covariates used in the Juvenile Production Index and Natal Rearing Proportional Passage models
<sup>2</sup>Covariates used only in the Natal Rearing Proportion Passage model

We designed two temperature metrics to represent both acute and chronic effects on incubating embryos. The first metric, average daily temperature from July 1 through September 30, covers the period when cold water volume in the reservoir is most likely limited and when most eggs hatch. This metric represents potential chronic effects of temperature on production across this period. The second metric, the number of degree-days over 12.0°C between May 1 and September 30, represents acute effects that occur once the threshold was exceeded. The threshold value was adopted by regulators as necessary to protect incubating eggs (NMFS 2024) based on an analysis by Martin et al. (2017).

A third temperature metric, mean temperature at RBDD (CDEC station RDB) during the peak month of juvenile winter-run catch (October), was designed to represent effects on juveniles rearing in the river prior to passage at RBDD. We assumed this temperature metric could reflect direct mortality attributed to temperature as well as indirect effects such as increased predator activity and disease susceptibility at warmer temperatures.

We considered five flow variables in total, though not all were included in every model. The first two were hypothesized to affect winter-run during egg incubation and emergence including, mean daily discharge at Bend Bridge (CDEC station BND; in cubic meters per second [m^3^/s]) during incubation and emergence, and total cumulative discharge (m^3^) at BND during incubation and emergence. The next three were hypothesized to affect non-natal and natal rearing life stages including, mean discharge (m^3^/s) at BND during the month of peak catch (or migration) at RBDD, the coefficient of variation in flow at BND during the month of peak catch (or migration) at RBDD, and mean daily maximum discharge (m^3^/s) at BND from September through November during emergence and migration.

### Data Analysis

Our analytical approach progresses systematically from drivers of total juvenile winter-run production regardless of migratory phenotype, to stock-recruitment relationships by migratory phenotype to test for density dependence and more nuanced models incorporating environmental drivers of life history expression. All analysis and data visualizations were carried out using R programming language within RStudio: Integrated Development Environment (R Core Team 2025; Posit team 2026).

#### Total Juvenile Production

We first examined how environmental conditions and spawner abundance influence total juvenile production. For this analysis, the response variable was the JPI which converts all life stages captured into “fry equivalents” as described earlier. We modeled the JPI as a function of female spawner abundance, environmental variables, and year to capture any variance in temporal trends for a total of 10 potential predictors. Prior to modeling, we examined correlations among variables and removed any highly correlated pairs (see supplemental material) and scaled the variables to standard z-scores.

Multiple flow and temperature variables could potentially influence variation in JPI. Thus, we further refined our list of potential factors using a Least Absolute Shrinkage and Selection Operator (LASSO) model to highlight important variables (McNeish 2015). Through regularization, LASSO models consider penalization for the number of included parameters within the modeling itself, as opposed to post hoc comparisons. Because of this, LASSO models perform well when the number of predictors exceeds the sample size or when strong correlations exist among predictors (Tibshirani 1996; Dormann et al. 2013). Unlike stepwise selection, LASSO models are not running multiple tests, and do not need Bonferroni corrections, and are less likely to be overfit due to “data dredging” than a stepwise procedure. Similarly, they are not sensitive to variable naming or order since everything is considered at once. Finally, LASSO model output contains quantitative measures of variable importance that are easy to interpret and extend to other analyses.

Based on the results of the LASSO model, we identified a set of candidate variables and used these to construct a series of *a priori* models. Priority was given to variables that managers can directly influence. These models were fitted using varying combinations of the candidate variables, alongside a null (intercept-only) model as a baseline for comparison. JPI is an integer count that is likely well modeled using a count process. Because the JPI data are over-dispersed, with variance significantly higher than the mean, we chose to model the process using negative binomial regression, with a square root link function to aid interpretability of effects.

Models were compared using Bayesian Information Criterion (BIC) scores to determine how well they were supported by the data while not being overfit. BIC was chosen over Akaike Information Criterion (AIC) due to the low number of data points and AIC’s propensity for overfitting with small sample sizes (Burnham and Anderson 2011). BIC model weights were used to select top candidate models and calculate model-averaged coefficients and unconditional confidence intervals for those models. Coefficients with unconditional confidence intervals that did not include zero were considered to have good support in the data (Burnham and Anderson 2011). Individual model predictive performance was assessed using leave-one-out cross-validation (LOOCV). For each observation *i*, we fit the model to all data excluding observation *i*, predicted the held-out value, and calculated the squared Pearson correlation between observed and predicted values across all folds (R^2^LOOCV).

Modeling results were tabulated to summarize model ranking metrics and model-averaged coefficients. Additionally, we constructed marginal effects plots for individual predictors using 95% confidence intervals to illustrate the predicted range of JPI across the observed values of each covariate. Finally, we used average model results to hindcast predicted JPI and overlay on observed JPI to further visualize predictive capability of the final model.

#### Proportion natal rearing model

This analysis addresses how environmental and demographic conditions influence the expression of migratory phenotype, specifically the proportion choosing to rear upstream for weeks to months (natal rearing) versus rapidly migrating downstream of RBDD (non-natal rearing). Annual proportion of natal to total non-natal passage was calculated as,

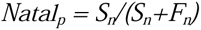

Where *Natal_p_*is the proportion of all juveniles rearing in natal habitat and then migrating, *S_n_* is the estimated natal rearing passage and *F_n_*is estimated non-natal rearing passage.

We again chose to use negative binomial regression due to overdispersed count data where we modeled natal rearing passage counts with log transformed total passage, sum of natal and non-natal rearing passage counts, as an offset variable. Because interpreting the offset this way requires a log link, we used a log-link function, in contrast to the square-root link used in the JPI model. Like the JPI model we used the same suite of covariates of female spawner abundance, environmental variables, and added one additional variable, *max_flow,* for a total of 10 potential predictors, all scaled by standard z-score. This approach preserves estimated natal rearing passage relative to total passage that would be lost by using proportions directly.

Variable selection followed the same LASSO regression approach described for the JPI model, using a Gaussian distribution and leave-one-out cross-validation given the small sample size (n = 23). Based on the results of the LASSO model, we identified a set of candidate variables and used these to construct a series of *a priori* models. Priority was given to variables that managers can directly influence. These models were fitted using varying combinations of the candidate variables, alongside a null (intercept-only) model as a baseline for comparison. Models were compared using BIC scores and R^2^ estimated from LOOCV. BIC model weights were used to select top candidate models and calculate model-averaged coefficients and unconditional confidence intervals for those models.

Modeling results were tabulated to summarize model ranking metrics and model coefficients, or model-averaged coefficients if applicable, with 95% CIs. To visualize these relationships, we generated marginal effects plots for individual predictors. All model predictions were generated with the passage offset set to 1, representing the predicted natal rearing proportion across the observed range of each covariate. Finally, we used the top-performing model to hindcast predicted natal rearing proportions, overlaying these values on observed data to evaluate the predictive capability of the final model.

#### Density dependence and juvenile production

We employed stock-recruit models to test for density dependent effects on total juvenile production and abundance of individual migratory phenotypes (non-natal and natal rearing). We hypothesized that because non-natal rearing individuals migrate soon after emergence and natal rearing individuals spend weeks or months in natal habitat before migrating, they can represent different mechanisms of density dependence. For example, a density-dependent relationship between spawners and non-natal rearing individuals may indicate limited spawning habitat availability whereas a relationship between spawners and natal rearing individuals may suggest that rearing habitat is limited. We first establish stock-recruitment dynamics with standard stock-recruit relationships to three response variables: total JPI, non-natal rearing passage, and natal rearing passage. For each, we compared a linear null model against models with Beverton-Holt and Ricker functional forms to characterize the underlying density-dependent processes. The linear model assumes a proportional relationship between spawner abundance and the production metric with no density dependence.

The Beverton-Holt model assumes strong density-dependent mortality at high densities, (Beverton and Holt 1993), and takes the following form,

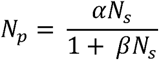

where *N_p_*is equal to the targeted production parameter, α is the productivity parameter, *N_s_* is spawner abundance, and *β* is the density dependence parameter.

The Ricker model assumes carrying capacity is approximated by the peak of the stock-recruit curve, with strong density-dependent mortality and overcompensation at high densities leading to reduced recruitment at very high spawner abundance (Ricker 1954),

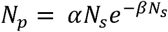

where *N_p_*is equal to the targeted production parameter, α is the productivity parameter, *N_s_* is spawner abundance, and *β* is the density dependence parameter.

We compared models using Bayesian Information Criterion BIC and selected the best-fitting model for each response variable. The model with the lowest BIC score was designated as the "best fit model" and the BIC value of this model was subtracted from the BIC values of other candidate models to calculate ΔBIC. We tabulated results for each individual modeling effort and visualized the results with smoothed stock-recruit curves overlayed over observed values.

## Results

Mean juvenile passage index (JPI) between 2002 and 2023 was 2.8 million (± 0.6 SE), ranging from 0.3 million (in 2022) to 8.9 million (in 2005). Non-natal rearing individuals comprised the majority of juvenile passage, averaging 2.0 million (± 0.5 SE) and ranging from 0.1 million (2022) to 7.5 million (2005). Mean natal rearing passage was 0.5 million (± 0.1 SE), ranging from 0.1 million (in 2022) to 1.0 million (in 2013). The proportion of natal rearing passage in relation to total juvenile passage averaged 0.28 (± 0.3 SE), ranging from 0.1 (in 2002) to 0.57 (in 2013). Summary statistics for environmental covariates are provided in Table 1, with additional details in the supplementary materials.

A detailed correlation matrix can be found in the supplementary material. The correlation analysis identified *Total_discharge_IE* and *Flow_IE* as the only variable pair exceeding our collinearity threshold (r > 0.99; Fig. S1), prompting removal of *Total_discharge_IE* from model fitting. *Temp_SAC_I* was highly correlated with *DD_above_12.0_I* (*r* = 0.92), *Flow_m* was highly correlated with *max_flow* (r = 0.92), and *Temp_SAC_I* was highly correlated with *Temp_RB_Oct* (*r* = 0.92); collinearity among these correlated variables was addressed during LASSO variable selection.

### Juvenile Production Index Modeling

The Lasso modeling process prioritized four key covariates: *Females*, *Flow_IE*, *Flow_CV_M*, and *DD_above_12.0_I*. Initial candidate models that included *year* showed it was not significant when combined with these covariates (p > 0.05), indicating that the measured environmental variables adequately explained temporal trends so *year* was excluded from all candidate models. Of the eight candidate models tested, the top three had substantial support (ΔBIC ≤ 1.5, cumulative weight = 0.89) and explained 59–73% of the variation in JPI (Table 2); the intercept-only model held very little weight (< 0.01), confirming that the included predictors meaningfully contribute to the model.

**Table 2.** Model selection results of the Juvenile Production Index negative binomial model. ΔBIC is based on model with lowest BIC, weight is the relative likelihood of the model based on BIC Weights, and R^2^ based on leave one out cross validation.

| Model | BIC | $\Delta BIC$ | weight | $R^2$ |
| --- | --- | --- | --- | --- |
| 1. Females+Flow_IE | 701.33 | 0 | 0.40 | 0.73 |
| 2. Females+Flow_IE+Flow_CV_M | 701.92 | 0.6 | 0.30 | 0.56 |
| 3. Females+Flow_IE+DD_above_12.0_I | 702.82 | 1.49 | 0.19 | 0.74 |
| 4. Females+Flow_IE+Flow_CV_M+DD_above_12.0_I | 703.96 | 2.63 | 0.11 | 0.56 |
| 5. Females+DD_above_12.0_I | 710.58 | 9.25 | <0.01 | 0.71 |
| 6. Females+Flow_CV_M+DD_above_12.0_I | 713.35 | 12.02 | <0.01 | 0.41 |
| 7. Females+Flow_CV_M | 718.12 | 16.79 | <0.01 | 0.26 |
| 8. Intercept Only | 732.22 | 30.89 | <0.01 | 0.00 |

Model averaged coefficients indicate that JPI is primarily driven by two factors: the number of female spawners (*Females*) and mean flow during the incubation and emergence period (*Flow_IE;* Table 3; Fig. 2). Both variables exhibited a direct, positive relationship with juvenile production. Other factors, such as flow variation at Red Bluff and water temperatures above 12°C at Highway 44, did not receive strong statistical support; evidenced by 95% CI overlapping 0 (Table 3). Overall, the model-averaged coefficients yielded predictions that closely mirrored observed JPI, with 2009 being the only notable outlier (Fig. 4A). These findings highlight a clear, strong link between demographic and hydrologic factors in driving annual juvenile recruitment.

**Table 3.** Model averaged parameters and unconditional 95% confidence intervals (standardized by *z*-score) for the JPI negative binomial model calculated using the four top models selected with BIC.

| Variable | Coefficient | CI_lower | CI_upper |
| --- | --- | --- | --- |
| Intercept | 1498.91 | 1341.59 | 1656.23 |
| Females | 526.74 | 332.77 | 720.71 |
| Flow_IE | 339.37 | 235.37 | 443.37 |
| Flow_CV_M | -62.16 | -246.72 | 122.4 |
| DD_above_12.0_I | -14.24 | -83.37 | 54.89 |

**Fig. 2.**
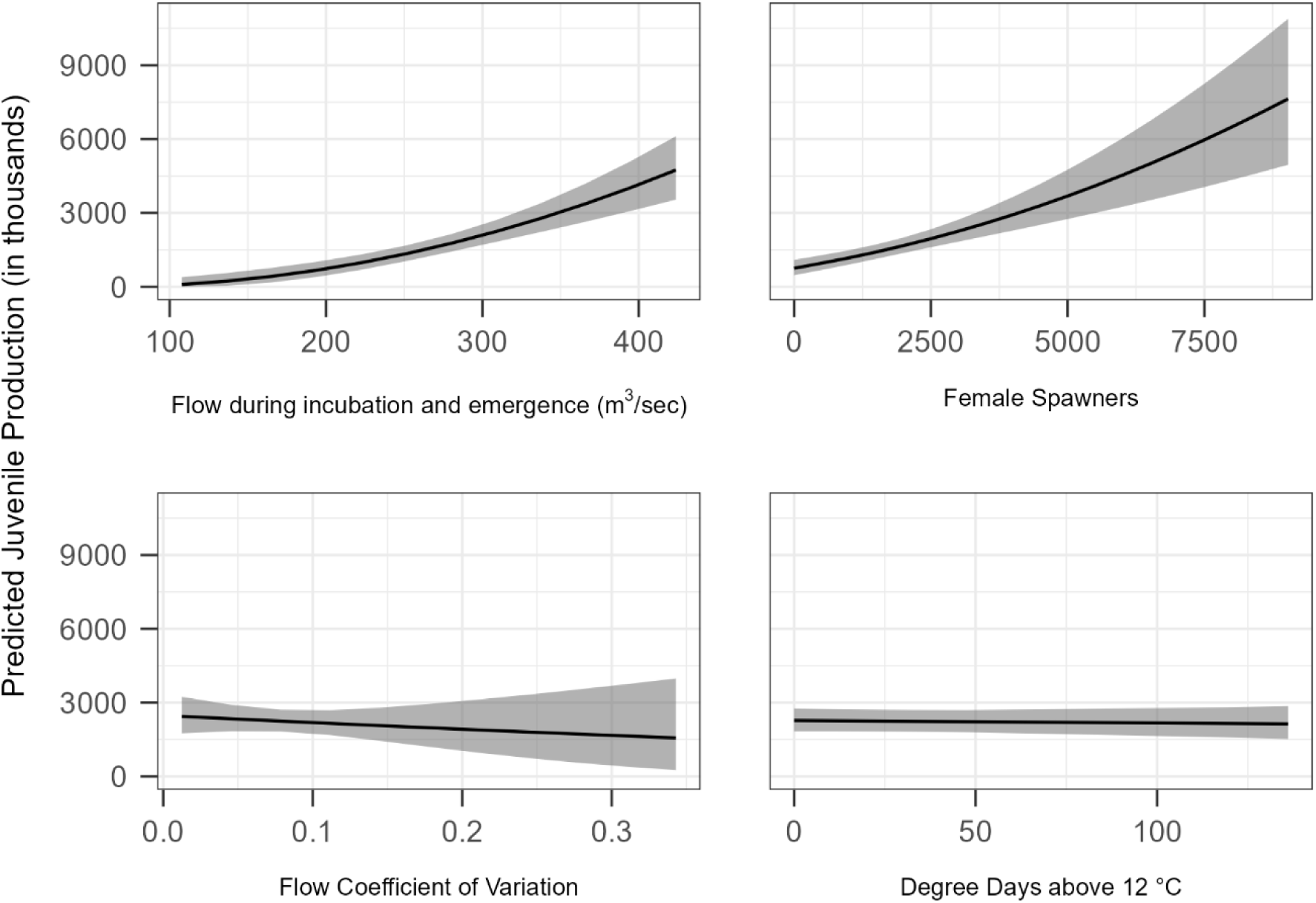
Marginal effects plots of model predictor variables from average model. Solid line represents predicted Juvenile Production Index and grey ribbon represents 95% confidence intervals.

### Proportion Natal Rearing Model

To investigate the environmental drivers of migratory phenotypes we modeled among-year variation in the proportion of natal rearing passage. While the Lasso model identified a similar suite of variables, including female abundance and various flow metrics, year remained a significant predictor in all candidate models (p < 0.05 in all cases), suggesting that important temporal trends were not fully captured by the measured environmental covariates. Year was therefore retained in all candidate models to control for unmeasured factors potentially influencing natal rearing migrant frequency. Of the seven candidate models tested, the top model had substantial support (ΔBIC ≤ 3.03, weight = 0.64; Table 5), and was the only candidate model carried forward for the remainder of the analysis. The intercept-only model held little BIC weight, confirming that the included predictors meaningfully contribute to the model. The top model captured similar variation (72%) as the JPI models, albeit with different predictor variables (Table 4).

**Table 4.** Model selection results of the proportion of natal rearers negative binomial model. Delta is based on model with lowest BIC, weight is the relative likelihood of the model based on BIC Weights, and R^2^ based on leave one out cross validation.

| Model | BIC | $\Delta BIC$ | Weight | $R^2$ |
| --- | --- | --- | --- | --- |
| 1. Year+Females+max_flow | 612.86 | 0.00 | 0.64 | 0.72 |
| 2. Year+Females+Flow_M | 615.89 | 3.03 | 0.14 | 0.71 |
| 3. Year+Females+Flow_IE+max_flow | 617.48 | 4.62 | 0.06 | 0.66 |
| 4. Year+Females+Flow_IE | 617.90 | 5.04 | 0.05 | 0.63 |
| 5. Year+Females+DD_above_12.0_I | 617.97 | 5.11 | 0.05 | 0.64 |
| 6. Year+Females+Flow_IE+max_flow+DD_above_12.0_I+Temp_RB_Oct | 619.00 | 6.14 | 0.03 | 0.71 |
| 7. Year+Females+DD_above_12.0_I+Temp_RB_Oct | 619.54 | 6.68 | 0.02 | 0.65 |
| 8. Intercept Only | 625.58 | 12.72 | 0.00 | 0.58 |

**Table 5.** Coefficients, standardized by *z*-score, and 95% confidence intervals for the proportion of natal rearers negative binomial model calculated using the top model for proportion of natal rearers.

| Variable | Coefficient | Lower 95% CI | Upper 95% CI |
| --- | --- | --- | --- |
| Intercept | -1.37 | -1.49 | -1.24 |
| Year | 0.27 | 0.12 | 0.42 |
| Females | -0.17 | -0.31 | -0.03 |
| max_flow | -0.18 | -0.32 | -0.04 |

After accounting for temporal trends (year effect: β = 0.27, p < 0.05), we found that proportion of natal rearing individuals was significantly and negatively associated with female spawner abundance (*Females*) and average daily maximum flow (*max_flow*; Fig 3; Table 5). The model accurately tracked interannual variability in proportional natal rearing passage; however, it substantially underestimated this proportion in 2012 and 2013, when observed rates reached their highest levels in the 23-year series and moderately overpredicted in 2022 and 2024 (Fig. 4B). Despite this, the results underscore that high flows and female spawner abundance are critical, albeit partial, predictors of the proportion of progeny rearing in natal habitat upstream of Red Bluff Diversion Dam.

**Fig. 3.**
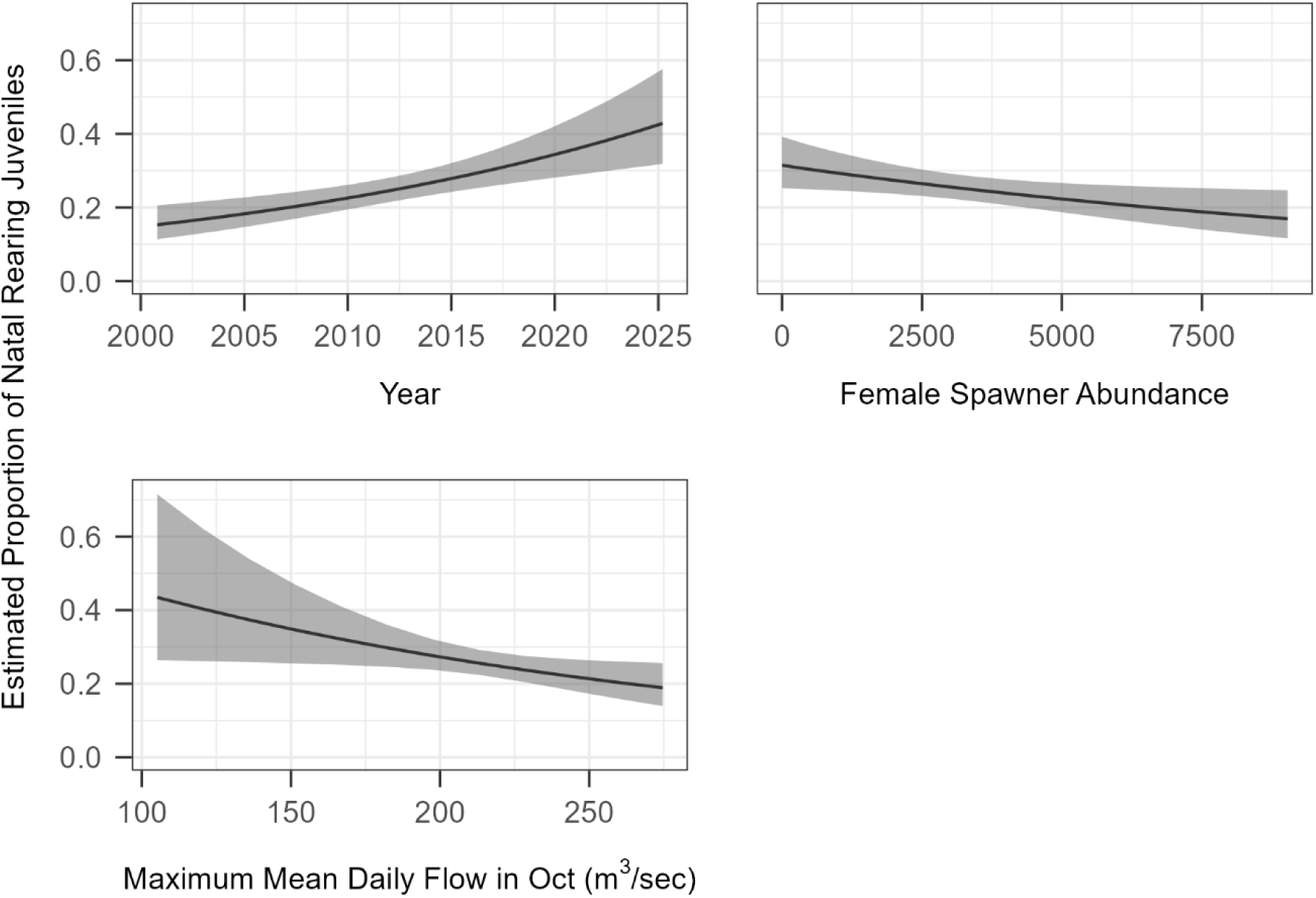
Marginal effects plots of model predictor variables from top candidate model for the proportional natal rearing model. Solid line represents predicted proportion of natal rearers and grey ribbon represents 95% confidence intervals.

**Fig. 4.**
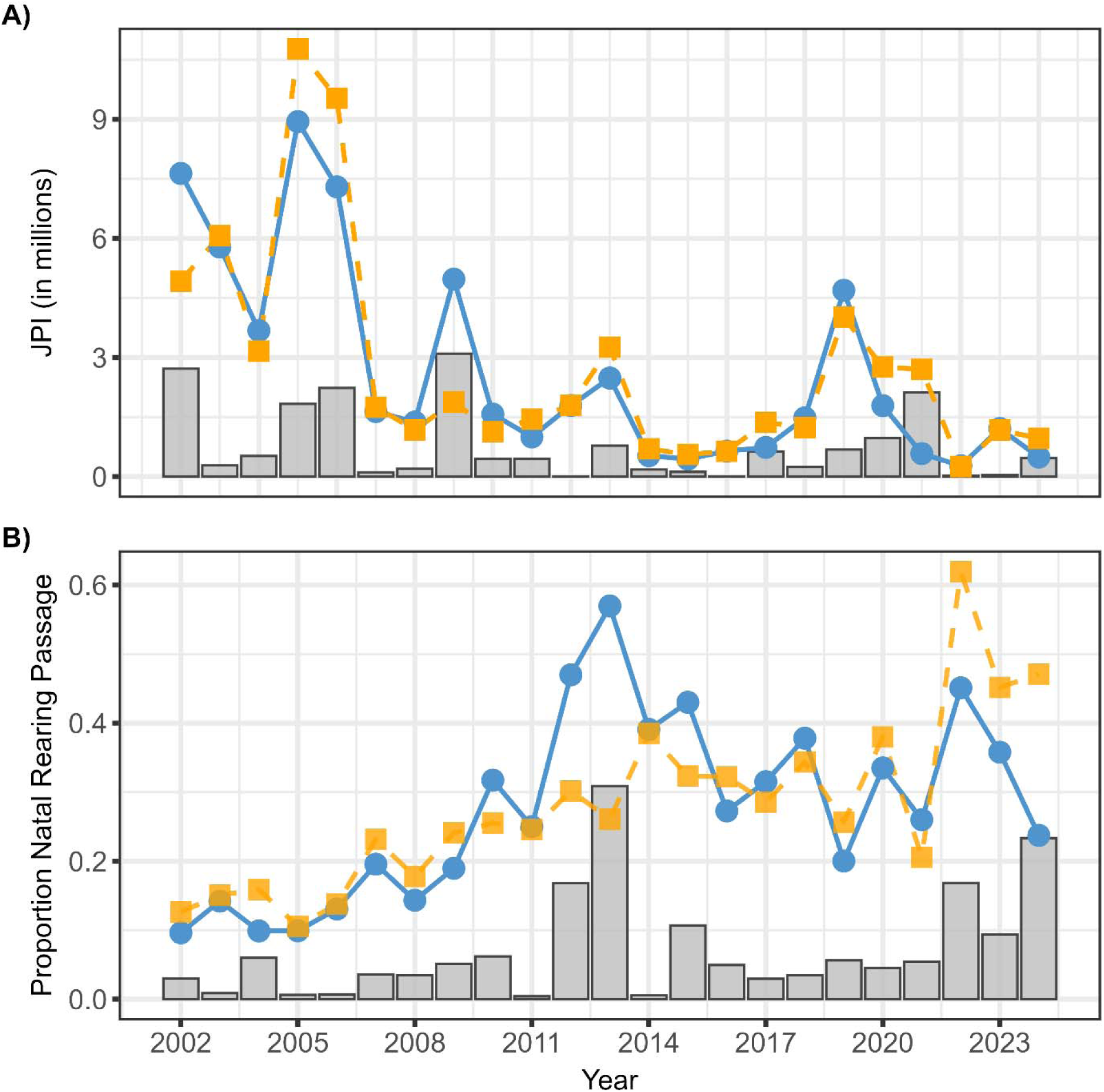
Predicted (dashed orange line and square points) and actual (solid blue line and circle points) values from the JPI negative binomial (A) and proportion of natal rearers negative binomal (B) models. Grey bars represent the difference between actual and predicted values.

### Stock-recruit relationships

Analysis using established stock-recruit relationships revealed that a linear model (density-independent) best described the relationship between female spawners and both total JPI and non-natal rearers based on BIC (Δ 3.06; Δ 2.14), with little support for Beverton-Holt or Ricker curves (Table 6; Fig. 5). However, a Beverton-Holt relationship (density-dependent) emerged as the best fit for the female-spawner – natal rearing passage relationship, with a Ricker model receiving modest support (Δ 0.79; Table 6; Fig. 5). Parameters for the best fit model (Beverton-Holt) indicated that in the absence of density-dependence, 361 natal rearers would be produced for each female spawner (α =361) and carrying capacity for natal rearers was estimated at 973,046 (α/β). These results suggest that while the non-natal life stage scales linearly with spawner abundance, the capacity for natal rearing is limited.

**Table 6.** Model summaries for stock-recruit models for Total Juvenile Production (Top), Non-natal Production (Middle), and Natal Rearing Production (Bottom).

| Model | df | Log likelihood | BIC | ΔBIC | Weight |
| --- | --- | --- | --- | --- | --- |
| *Total Juvenile Production* |  |  |  |  |  |
| Linear (Density independent) | 2 | -360.24 | 726.75 | 0.00 | 0.70 |
| Beverton-Holt | 3 | -360.20 | 729.81 | 3.06 | 0.15 |
| Ricker | 3 | -360.21 | 729.82 | 3.07 | 0.15 |
| *Non-natal Rearing Production* |  |  |  |  |  |
| Linear (Density independent) | 2 | -358.20 | 722.67 | 0.00 | 0.59 |
| Beverton-Holt | 3 | -357.70 | 724.81 | 2.14 | 0.20 |
| Ricker | 3 | -357.71 | 724.83 | 2.16 | 0.20 |
| *Natal Rearing Production* |  |  |  |  |  |
| Beverton-Holt | 3 | -317.12 | 643.65 | 0.00 | 0.55 |
| Ricker | 3 | -317.52 | 644.44 | 0.79 | 0.37 |
| Linear (Density independent) | 2 | -320.67 | 647.62 | 3.97 | 0.08 |

**Fig. 5.**
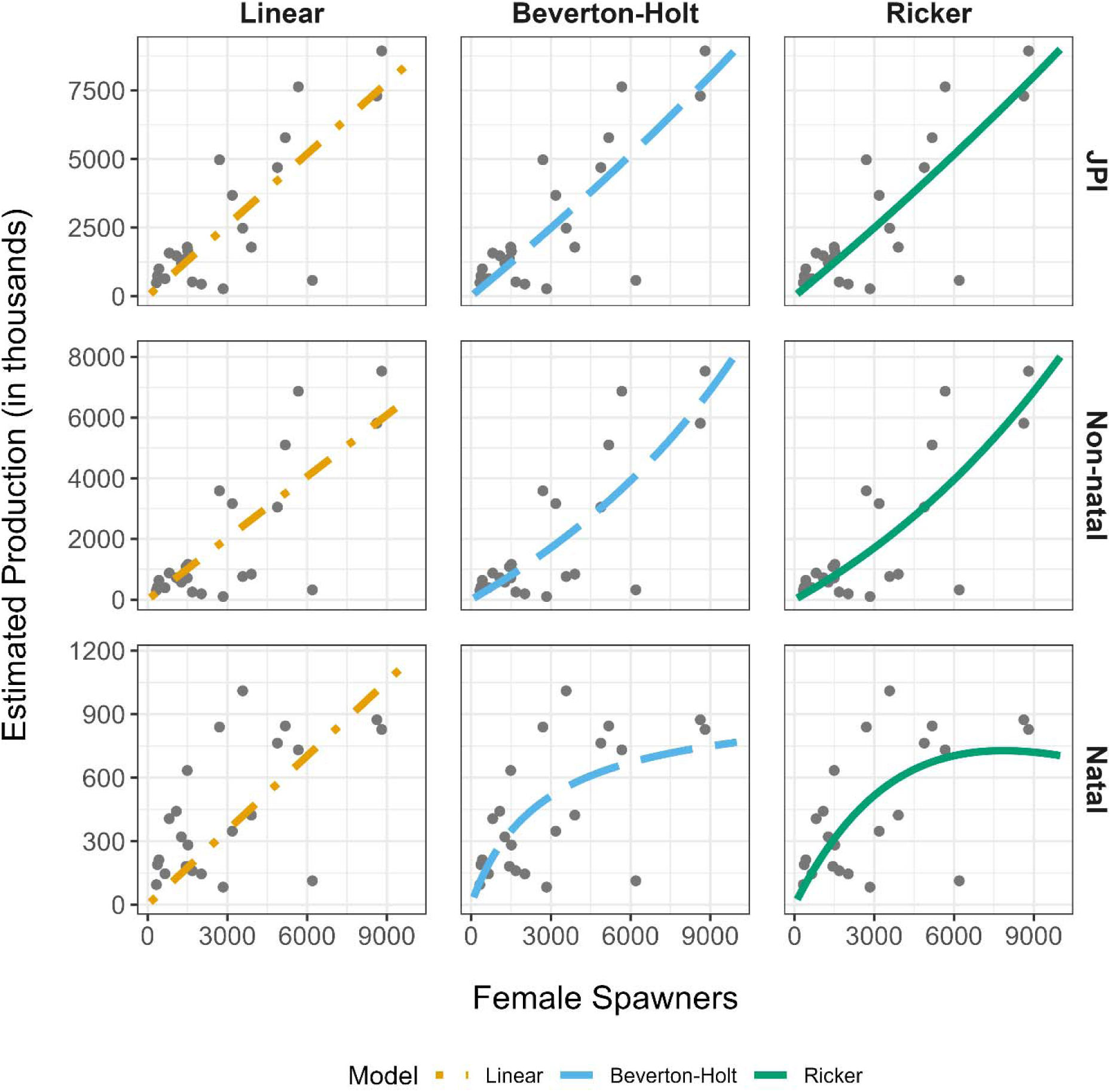
Graphs of fitted stock-recruit models for each population faceted by stock-recruit models (columns) and life stage (rows). Linear models are represented by dot-dash orange lines, Beverton-Holt models by long-dash blue lines, and Ricker models by solid green lines. JPI refers to Juvenile Production Index, and non-natal and natal refer to non-natal rearing and natal rearing passage estimates respectively as described in the text.

## Discussion

Our analysis of more than two decades of monitoring data revealed population dynamics and environmental relationships that can directly inform management and restoration actions for endangered winter-run. After accounting for spawner abundance, our results highlight river flow during egg incubation and fry emergence as a strong driver of juvenile production. However, the modeled relationships reported here cannot isolate the specific mechanisms driving this flow benefit. In salmonid early life stages, the impact of flow varies depending on the interaction between habitat structure and flow magnitude (Arthaud et al. 2010; Armstrong and Nislow 2012). During incubation, hyporheic water flow is crucial for delivering oxygen to the redd and removing metabolic waste (Johnson 1980; Lapointe et al. 2004); conversely, excessively high flows can mobilize substrate and scour eggs from the gravel (May et al. 2009; Gendaszek et al. 2018). Following emergence, flow continues to affect fry and parr by altering rearing habitat quality, food delivery rates, and migratory cues (Harvey et al. 2006; Zeug et al. 2014; Sturrock et al. 2015; Warkentin et al. 2022). When interpreting these mechanisms within our study, it is important to note that flows during winter-run incubation and emergence are highly regulated with observed mean daily flow ± standard deviation being higher but significantly less variable (244 ±70 m^3^/s) from May-October compared to the spectrum of flows observed across the full year in the study area (197 ±250 m^3^/s; see Supplementary Material Fig. S4). Consequently, extreme low flows (which dewater incubating eggs) and extreme high flows (which cause scouring) are generally avoided.

It was unexpected that temperature did not have a stronger influence on juvenile production because thermal stress during egg incubation has been implicated as the primary factor limiting winter-run early life stage survival (Martin et al. 2017; Anderson et al. 2022). These previous analyses used the JPI and female spawner abundance data to calibrate mechanistic models for temperature mortality potentially occurring during egg incubation. Generally, they found considerable interannual variation in estimated egg-to-fry survival could be explained by water temperatures experienced during egg incubation. However, neither Martin et al. (2017) nor (Anderson et al. (2022) evaluated alternative models that included flow as a covariate. Our findings, however, suggest temperature may not be the sole contributor to mortality during incubation. For example, Martin et al. (2020) found that thermal tolerance of embryos depends on complex, nonlinear interactions between developmental stage, ambient dissolved oxygen concentration, flow velocity, and the presence of neighboring eggs. Notably, they found that egg survival was poor at low water velocities regardless of incubation temperature, suggesting that adequate flows are essential even when thermal targets are met. However, the modeled flow response may reflect post-emergence processes rather than strictly egg survival. For example, they can influence observed production by cuing and facilitating dispersal and downstream migration of fry, a pattern that is broadly supported by many previous Central Valley salmon investigations (e.g. Sommer et al. 2005; Del Rosario et al. 2013; Zeug et al. 2014; Munsch et al. 2020; Sturrock et al. 2020; Burford et al. 2025). However, flow events that might cue fry migration are absent during the emergence period (September – October). Higher magnitude flow years may simply increase displacement of fry or improve survival of fry moving between spawning areas and RBDD.

Our results do not imply that temperature management is unimportant. Because winter-run Chinook salmon currently spawn far downstream of their historic, high-elevation habitat, maintaining cold water via the Temperature Control Device (TCD) remains critical to avoiding significant early life-history impacts (USFWS 1999). Even under the current management regime, severe drought years can deplete the cold-water pool (Hallnan et al. 2020), risking a loss of thermal control just as fry begin to emerge. However, because operations tightly optimize water temperatures to meet strict regulatory targets, the day-to-day and year-to-year operational variation is effectively shifted onto discharge. Consequently, by successfully minimizing thermal stress, the management regime may ensure that flow-mediated dynamics and spawner abundance determine a large fraction of the year-to-year variation in juvenile production.

There was strong evidence that migratory phenotypes responded differently to environmental drivers. In most years, non-natal rearing accounted for the largest fraction of juvenile migrants. Yet as maximum discharge increased, the proportion of non-natal rearing migrants also increased. Zeug et al. (2014) reported a similar pattern for fall run Chinook Salmon where both flow magnitude and flow variance increased the proportion of juveniles migrating as fry. Increasing flow magnitude may physically displace newly emerged fry from natal habitat as their swimming ability is still limited (Saltveit et al. 1995). In addition, Tiffan et al. (2002) found a general decrease in available rearing habitat as flows increase, suggesting that the lack of suitable rearing habitat for early life stages may induce downstream dispersal. Regardless of the specific mechanism, Phillis et al. (2018) found that up to 65% of winter-run Chinook Salmon adults reared as juveniles in tributaries to the Sacramento River or in the Delta. Juvenile winter-run entered these habitats at small sizes, but departed freshwater at a similar size as those reared in the mainstem Sacramento River. Consequently, juvenile salmonids that migrate downstream and rear in non-natal habitats have been shown to make substantial contributions to subsequent adult spawning escapement in multiple river systems (Koski 2009; Miller et al. 2010; Bennett et al. 2015; Sturrock et al. 2015), suggesting that higher discharge could be facilitating access to these non-natal rearing habitats and potentially increasing the success of this life history strategy.

Our findings showed that the effect of female spawner abundance also diverged among migratory phenotypes; a well-supported density-dependent relationship was evident for natal rearing fish whereas the relationship with non-natal rearing fish was linear. This indicates that the amount of suitable habitat for juvenile rearing is a limiting factor, at least for the natal rearing component of the population. While the aggregated JPI metric serves as a valuable "common currency" within regulatory frameworks to estimate total juvenile production, the stock-recruit modeling reported here suggests it may obscure important population dynamics. Otolith strontium isotope analysis showed that 34% to 56% of juvenile winter-run had reared primarily in the mainstem Sacramento River (Phillis et al. 2018). However, this otolith analysis could not identify where along the 412 rkm Sacramento River this extended rearing had occurred. Mean natal rearing passage estimates reported here (0.5 million ± 0.1 SE) suggest a substantial fraction of mainstem Sacramento River rearing identified by Phillis et al. (2018) may have occurred upstream of Red Bluff Diversion Dam. Offsetting the limited capacity of natal rearing habitat, juvenile rearing habitats at or near spawning areas are expected to provide consistently favorable environmental conditions, and potentially reduced exposure to predation, as evidenced by the low outmigration survival in downstream habitats observed across multiple acoustic telemetry studies (e.g. Michel et al. 2015; Hassrick et al. 2022). In contrast, downstream non-natal habitats are likely more stressful in drought years, but providing better opportunities for growth and survival primarily during cooler, wetter seasons (Coleman et al. 2022). Despite downstream risks, however, phenotypes that are less numerically abundant have been found to contribute disproportionately to the adult population (Cordoleani et al. 2021). Consequently, evaluating juvenile production purely through a single, aggregated index may overlook the life-history diversity that can buffer the population against environmental stochasticity and obscure factors limiting juvenile production.

The mechanisms controlling density dependence are not known but some inference can be made. The lack of density dependence between female spawners non-natal rearing fish suggests that spawning habitat quantity is not limiting at current population levels as redd superimposition (and thus embryo mortality) would be expected at high spawner densities (Dudley 2019). However, this does not necessarily reflect the quality of spawning habitat— improvements in habitat productivity could shift the intercept of the linear relationship higher (more recruits-per-female). Einum et al. (2006) found that the mechanisms of density dependence in juvenile salmonids can change throughout ontogeny as the ability to respond to stressors changes. Non-natal rearing winter-run juveniles in this analysis would have begun migrating shortly after emergence; a well-documented life-history strategy among

Central Valley salmonids (Miller et al. 2010; Sturrock et al. 2015; Phillis et al. 2018). This suggests an ability to disperse in response to localized density-dependent pressures as fish transition to different life stages (Sorel et al. 2023). However, habitat for these dispersing fish appears limited via density dependence. Consequently, restoration actions targeted at improving rearing habitat capacity would be expected to yield higher per-capita production across all spawner densities. For example, the steady increase in natal rearing passage coincides with concerted restoration efforts. These projects improved side-channel connectivity, flow, and depth in marginal rearing zones, while adding rock wads to deepwater habitats and injecting gravel to expand spawning areas (Banet et al. 2023; U.S. Bureau of Reclamation 2024). While we cannot definitively attribute this increase in natal rearing proportion to restoration efforts without direct evaluation, if improved habitat availability is contributing to this trend, it would suggest that expanding upstream rearing capacity may promote an increase in natal rearing.

Recognizing the value of multiple life-history pathways, the challenge lies in identifying a scheme to support these strategies that facilitates population recovery. Our results suggest that a combination of flow management and habitat restoration, while maintaining a suitable thermal regime, will be needed. Sturrock et al. (2020) found that the stabilized, anthropogenically modified flow regimes in the Central Valley may be selecting against non-natal rearing and reducing the population’s life history plasticity. Implementing a ‘facilitated migration’ framework as demonstrated by Burford et al. (2025), may offer a partial path forward. This approach focuses on strategic, pulse-mediated flow releases designed to align reservoir operations with natural migration phenology. By doing so, managers could not only increase the overall abundance of oceanward migrants, but also actively promote diverse downriver rearing strategies (Zeug et al. 2014; Sturrock et al. 2015) provided that fish are moved downstream when appropriate rearing habitats are activated. However, any flow strategy will have limited effectiveness if there are physical habitat limitations on population productivity, such as the density-dependent relationship between female spawners and natal rearing we described here. Failure to address habitat limitation while implementing targeted migration flows may increase per-capita survival during migration only to lose those gains to density dependence during subsequent life stages.

This analysis highlights the need for a diversified portfolio of management actions that extends beyond maximizing individual metrics to optimize the full suite of environmental conditions that support all life stages. This is especially important as climate change is projected to further compress the operational bandwidth by warming temperatures and reducing dissolved oxygen, complicating the already narrow window for flow and temperature management. Effective conservation of winter-run Chinook Salmon requires integrating: (1) optimized flow management to maintain suitable conditions during egg incubation and fry emergence, (2) strategic habitat restoration to expand rearing capacity upstream of RBDD, (3) operational flexibility that balances natal rearing production with downstream non-natal rearing by timing flow releases to coincide with habitat activation, and (4) continued monitoring to evaluate population-level responses to these integrated actions and support adaptive management.

## ACKNOWLEDGEMENTS

We thank the Sacramento River Settlement Contractors and the United States Bureau of Reclamation for funding this study. We are grateful to the California Department of Fish and Wildlife and the United States Fish and Wildlife Service for the collection and curation of the data analyzed here.

## AUTHOR CONTRIBUTION

All authors contribution to the study conception, design, and writing. Initial conceptualization and analyses were conducted by Cavallo, Ross, and Zeug. Final modeling and first draft of the manuscript was written by Ehlo with input provided by all authors before approving and submitting the final manuscript

## DATA AVAILABILITY

The summarized dataset used for the analysis and the analysis code are available upon request. Raw data used in the analysis can be queried from multiple sources including Central Valley Prediction and Assessment of Salmon https://www.cbr.washington.edu/sacramento/, Environmental Data Initiative https://edirepository.org/, CalFish https://www.calfish.org/Home.aspx, and California Data Exchange Center https://cdec.water.ca.gov/.

## ETHICS STATEMENT

This study used data from the U.S. Fish and Wildlife Service Red Bluff Diversion Dam Rotary Screw Trap monitoring and the California Department of Fish and Wildlife Chinook Salmon carcass surveys. Both surveys follow established state and federal protocols for the monitoring of Chinook Salmon in the Central Valley of California. As the analyses were conducted solely on existing data collected under these standardized procedures, no additional ethical approval was required.

