## Supplemental Materials for "Winter-run Chinook salmon juvenile recruitment and early life history response to flow in a heavily altered tailwater"

**Table S1.** Environmental variables considered for modeling both JPI and smolt proportion. Description of the metric, temporal variation, and hypothesized effects are included. All data were queried from California Data Exchange Center (CDEC). <https://cdec.water.ca.gov/dynamicapp/staSearch>

| **Covariate Name** | **Flow metrics description** | **Months** | **Hypothesis** |
| --- | --- | --- | --- |
| Flow_IE^1^ | Mean flow during incubation and emergence (m^3^/sec) at Bend Bridge (BND) | May-October | Mortality effects of flow occur at daily time steps during incubation and emergence |
| Total_discarge_IE^1^ | Cumulative discharge (m^3^) during incubation and emergence at BND | May-October | Mortality effects of flow occur at a seasonal time step in response to the total volume of water discharged during incubation and emergence |
| Flow_M^1^ | Mean flow during month of peak migration (m^3^/sec) at BND | October | Survival during migration is directly proportional to flow magnitude |
| Flow_CV_M^1^ | CV of flow during peak month of migration at BND | October | Flow pulses increase survival (Hassrick et al. 2022) |
| max_flow^1^ | Maximum daily flow (m^3^/sec) averaged by year across the emergence and migration period at BND | October | Hydro peaking may initiate early outmigration of winter-run fry |
| Temp_SAC_I^2^ | Mean water temperature (°C) during key incubation period (Jul-Sep) at Hwy 44 (SAC). Indexes average temperature experienced by incubating eggs | July-September | Temperature effects on survival are primarily chronic and increase linearly with mean daily values in the period when cold water volume is limiting. |
| DD_above_11.67_I^2^ | Cumulative degrees per day above 11.67°C during incubation period at Hwy SAC. Indexes magnitude of exposure to temperature stress | July-September | Temperature effects on production are primarily acute and occur when a threshold value is exceeded |
| Temp_RB_M^3^ | Mean water temperature (°C) at Red Bluff (RDB) during month of peak migration. Indexes metabolic demand of juveniles and their predators | October | Higher temperatures during rearing/migration increase predation intensity and demand for prey |
| ^1^https://cdec.water.ca.gov/dynamicapp/staMeta?station_id=BND  ^2^https://cdec.water.ca.gov/dynamicapp/staMeta?station_id=SAC  ^3^https://cdec.water.ca.gov/dynamicapp/staMeta?station_id=RDB | | | |

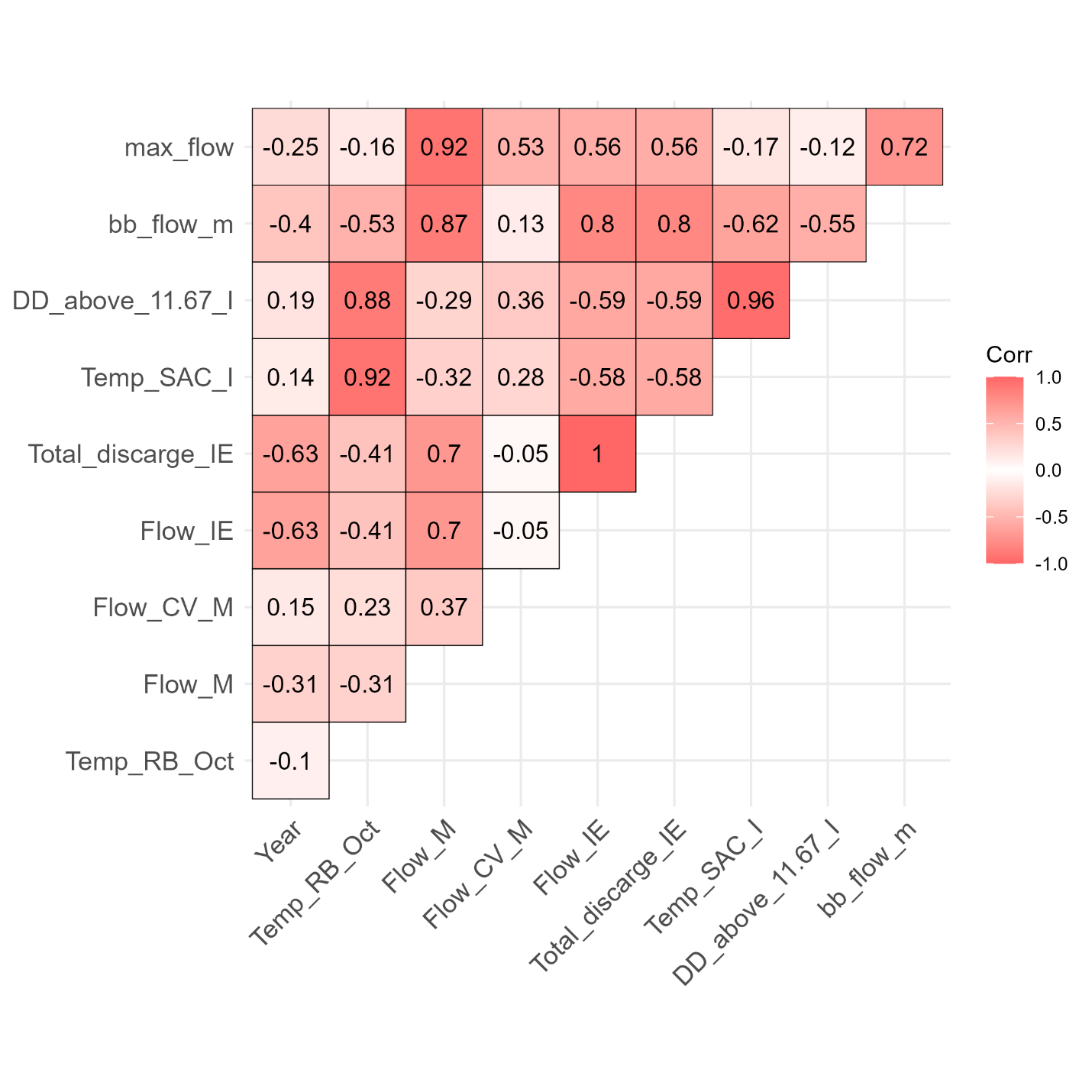

Figure S1. Correlation matrix of variables used in modeling for both JPI and proportion of smolts. Numbers closer to 1 or -1 indicate greater positive or negative collinearity.

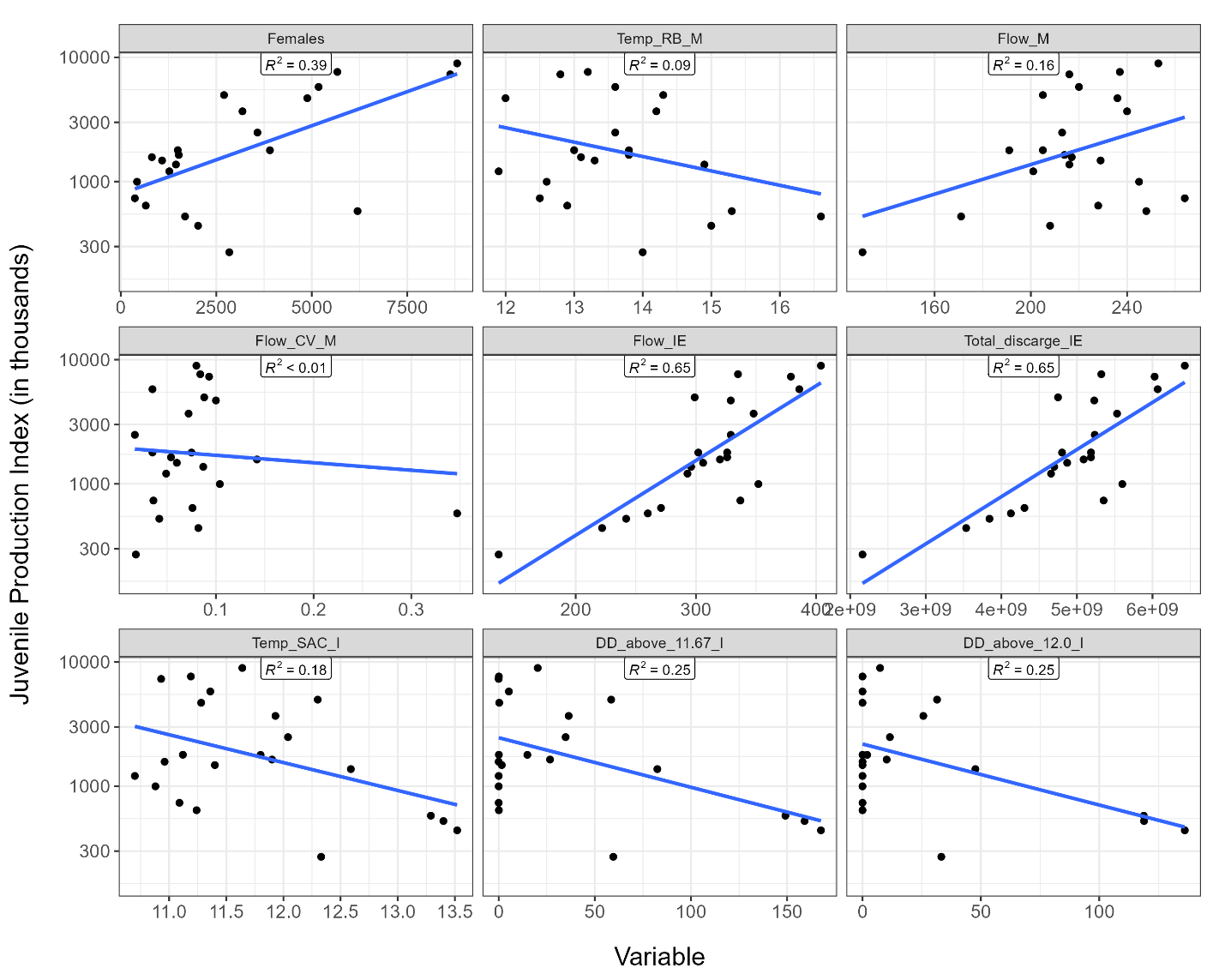

Figure S2. Exploratory simple linear regression analysis of environmental variables and the Juvenile Production Index. R^2^ values represent coefficients of determination and solid blue line indicates fitted linear regression. Note that the y-axis is presented on the logarithmic scale.

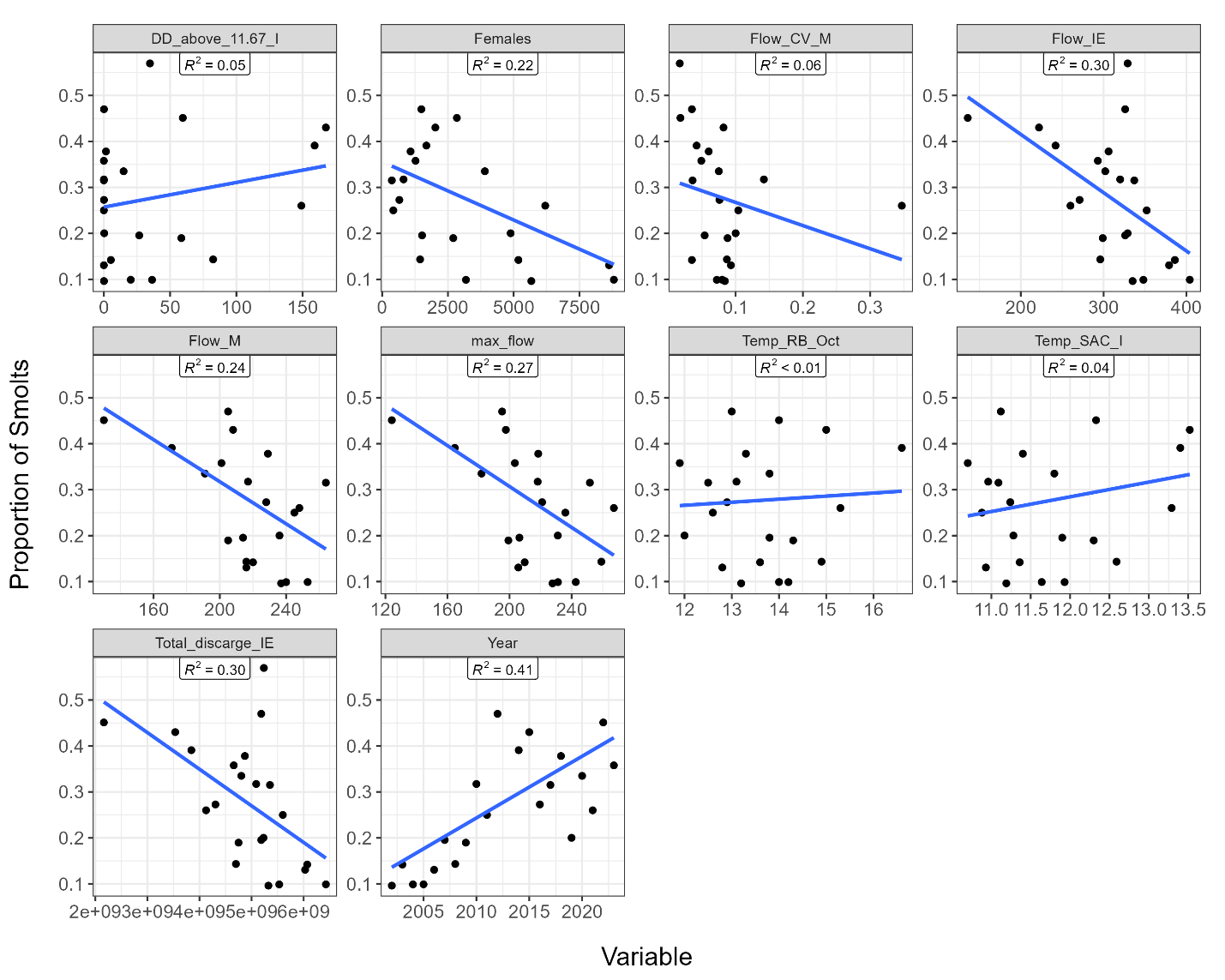

Figure S3. Exploratory simple linear analysis of environmental variables and proportion of smolts. R^2^ values represent coefficients of determination, and solid blue line indicates fitted linear regression.

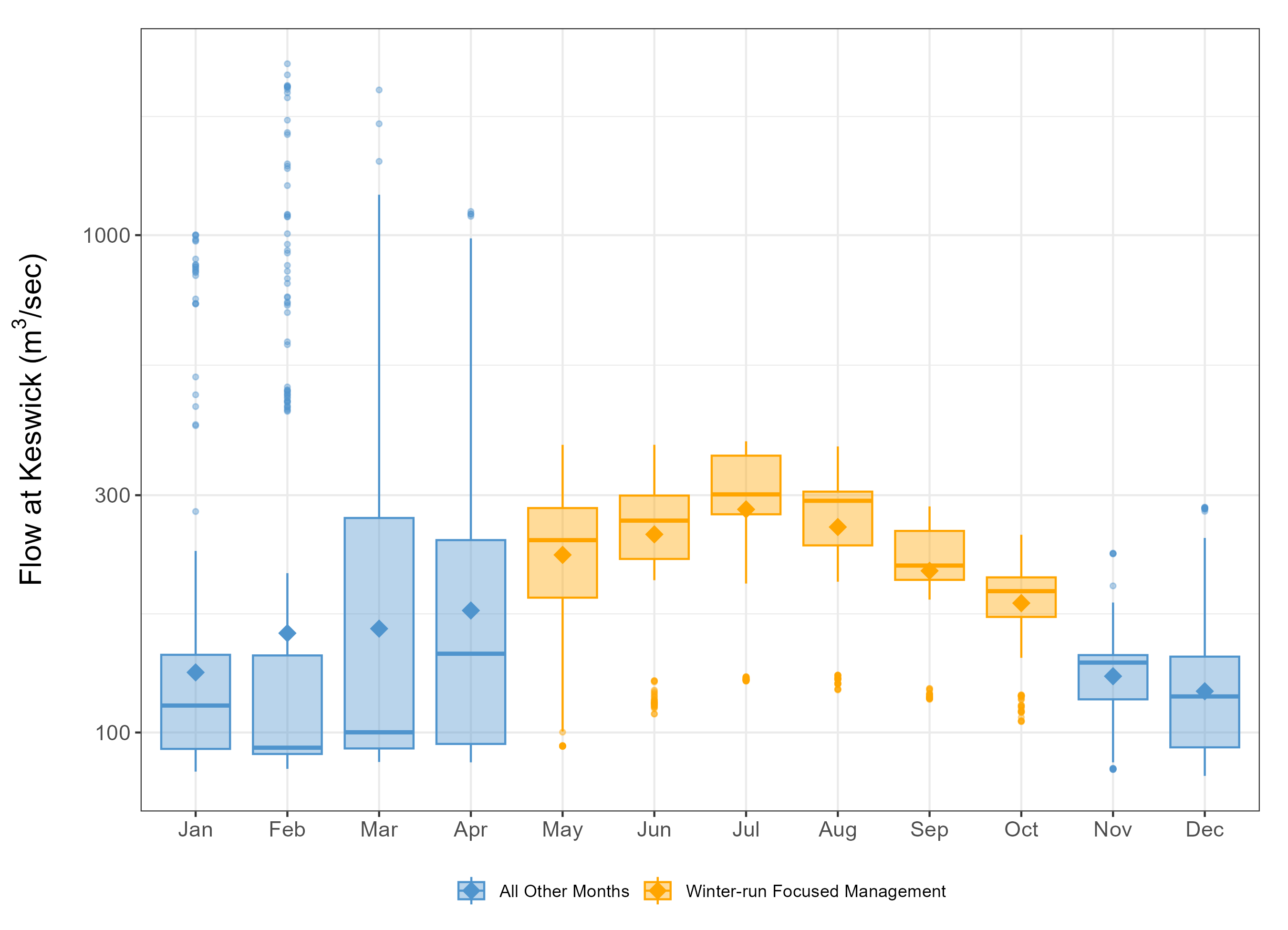

Figure S4. Boxplot summary of daily flow releases from Keswick Dam into the Sacramento River from 2002-2024. Diamonds represent average flow by month, orange represents months in which releases are focused around temperature and flow management for winter-run spawning, incubation, and migration, and blue represents months in which flow management is subject to other controlling factors. Note that the y-axis for flow is on a logarithmic scale.
